# DIPA-CRISPR gene editing in the yellow fever mosquito *Aedes aegypti* (Diptera: Culicidae)

**DOI:** 10.1101/2023.04.07.535996

**Authors:** Yu Shirai, Momoyo Takahashi, Manabu Ote, Hirotaka Kanuka, Takaaki Daimon

**Affiliations:** Department of Applied Biosciences, Graduate School of Agriculture, Kyoto University, Kitashirakawa Oiwakecho, Sakyo-ku, Kyoto 606-8502, Japan; Department of Tropical Medicine, The Jikei University School of Medicine, Tokyo, Japan; Center for Medical Entomology, The Jikei University School of Medicine, Tokyo, Japan

**Keywords:** DIPA-CRISPR, mosquito, vitellogenesis, ovary, *kynurenine 3-monooxygenase*

## Abstract

Current methods for gene editing in insects rely on embryonic microinjection, which can be challenging for non-specialist laboratories. Recently, an alternative method known as “direct parental” CRISPR (DIPA-CRISPR) was developed. This method involves injecting commercial Cas9 protein and single-guide RNA into adult females, which can efficiently introduce mutations into developing oocytes. However, its versatility has not been fully explored, particularly in insects that have the most derived, polytrophic meroistic ovaries. In this study, we successfully applied DIPA-CRISPR to the yellow fever mosquito *Aedes aegypti*, which has polytrophic meroistic ovaries. Following adult injection of Cas9 ribonucleoproteins (Cas9 RNPs) targeting the kynurenine 3-monooxygenase gene, we recovered gene-edited G_0_ individuals. Injection at 24 h after blood-feeding resulted in the highest gene editing efficiency (3.5%), confirming that a key parameter of DIPA-CRISPR is the stage in which the adult females are injected. Together with our previous study, we demonstrated that DIPA-CRISPR is applicable to all three types of insect ovaries (i.e., panoistic, telotrophic, and polytrophic), which indicates that DIPA-CRISPR is a generalizable approach for insect gene editing.

## Introduction

The development of genome editing tools has enabled sophisticated genome engineering in insects (Gantz and Akbari 2018; Matthews and Vosshall 2020). Current methods for gene editing in insects rely on embryonic microinjection at the preblastoderm stage, which requires expensive equipment and high technical skill. These limit the application of gene editing methods to a wide variety of insect species. For example, microinjection into an early embryo is very difficult or virtually impossible in species that lay their eggs inside their prey (e.g., parasitoid wasps), produce live larvae instead of eggs (e.g., viviparous aphids and flesh flies), or produce a hard egg case encapsulating their eggs (e.g., cockroaches).

Recently, an alternative method known as “direct parental” CRISPR (DIPA-CRISPR) was developed, which enables gene editing by simple adult injection (Shirai et al. 2022). This technique only requires two components: commercial Cas9 protein and single-guide RNA (sgRNA). Cas9 ribonucleoproteins (RNPs) injected into adult females are incorporated into developing oocytes from the hemolymph along with yolk protein precursors, thereby introducing heritable mutations into the oocytes. We demonstrated previously that DIPA-CRISPR enables highly efficient gene editing in the German cockroach *Blattella germanica* and the red flour beetle *Tribolium castaneum* (Shirai et al. 2022).

Despite its simplicity, the applicability of DIPA-CRISPR to a wider range of insects has not been fully explored. There are three distinct organizations of insect ovaries, which include panoistic, telotrophic meroistic, and polytrophic meroistic ovaries (Mclaughlin and Bratu 2015). Although we showed that DIPA-CRISPR is applicable to insects with panositic (*B. germanica*) and telotrophic (*T. castaneum*) ovaries, it is unclear whether it is also applicable to insects with the most derived, polytrophic meroistic ovaries. In this study, we evaluated DIPA-CRISPR in the yellow fever mosquito *Aedes aegypti* (L.) (Diptera: Culicidae), which has polytrophic ovaries. We provide evidence that DIPA-CRISPR works in *A. aegypti*, suggesting that this simple, accessible method can be applied to insects with all three types of ovaries.

## Materials and Methods

### Insects

The *A. aegypti* strain was derived from the Liverpool strain and was kindly provided by Dr. Ryuichiro Maeda (Obihiro University of Agriculture and Veterinary Medicine). Eggs were obtained by a standard procedure and incubated in reverse osmosis (RO) water. The hatched first instar larvae were transferred to a plastic container and provided RO water and daily fish food (Hikari, #4971618, Kyorin Co., Ltd., Japan). The larvae were maintained in an insectary room at 27 °C, and the water was refreshed every 2–3 days. Pupae were collected in a plastic cup and placed in a cage (bottom 27 cm × 27 cm, top 25 cm × 25 cm, height 27 cm) with a 50-ml glass flask containing 10% sucrose solution with an inserted filter paper (#1001-125, Whatman, USA). The cage in which the emerged adults were reared was maintained in an incubator (MIR254-PJ, Panasonic Co., Japan) at 27 °C with humidity over 90% in a standard 12 h:12 h light:dark cycle. The sucrose solution was changed every 3–4 days. For injection experiments, the adult females were fed mouse blood and separated from males prior to injection.

### Preparation of Cas9-sgRNA RNPs

sgRNAs targeting *A. aegypti kmo* (GenBank: NC_035108.1) were synthesized as previously described (Shirai and Daimon, 2020; Shirai et al. 2022). Briefly, annealed oligo DNA was cloned into the *BsaI* site of the pDR274 vector (Hwang et al., 2013). After linearization with *DraI*, the vector was used as a template for in vitro transcription using the T7 RiboMAX Express Large Scale RNA Production System (Promega, Cat#P1320). The synthesized sgRNAs were extracted with phenol (pH 4–5):chloroform:isoamyl alcohol (125:24:1) (Sigma, Cat#P77619), precipitated with isopropanol, and dissolved in RNase-free water. Commercial Cas9 protein was purchased from IDT (Alt-R S.p. Cas9 Nuclease V3, Cat#1081059). Cas9 protein and sgRNAs were mixed at a molar ratio of approximately 1:2 and incubated for 10–15 min at room temperature to allow Cas9 RNP formation. The target sequences of the sgRNAs are (5’- to -3’): AAGACCAGGCCTCAATCGT for sgRNA1 and CGGCAAGGCGGTGATCAT for sgRNA2.

### Adult injections and screening

Injections were performed using a glass capillary needle equipped with an IM 300 Microinjector (NARISHIGE). Females were anesthetized on ice. Approximately 200 nL of the Cas9 RNP solution containing 3.3 μg/μL Cas9 (IDT, Cat#1081059) and 1.3 μg/μL sgRNAs (a mixture of sgRNA1 and sgRNA2) was injected into the thorax of the adult females. The females were subsequently grouped in a cage (bottom 15 cm × 15 cm, top 15 cm × 15 cm, height 15 cm). The eggs were laid onto filter paper soaked in RO water beginning 2 days after blood-feeding. The G_0_ progeny were collected for 3–4 days after the paper was set. The eye colors of the G_0_ generation were examined at the pupal or adult stage, and some individuals were subjected to individual genotyping.

### Genotyping

Genomic DNAs were extracted individually as previously described (Daimon et al., 2014). Genomic PCR was performed using KOD FX Neo (TOYOBO, Cat#KFX-201). Mutations were screened by analyzing the PCR products from a heteroduplex mobility assay (HMA) using the MultiNA Microchip Electrophoresis System (MCE-202, Shimadzu). Primer sequences for HMA are (5’- to -3’): ATTGGTCGTGAGCGGTTGG and GTACAATCCTCGAATCCGGCATTC. Primer sequences for Sanger sequencing are (5’- to -3’): GCACTTGGACGGTGACGCTG and GTACAATCCTCGAATCCGGCATTC.

## Results

### Target gene for DIPA-CRISPR in *A. aegypti*

To evaluate the efficiency of DIPA-CRISPR in *A. aegypti*, we targeted the kynurenine 3-monooxygenase (*kmo*) gene, which is essential for eye pigmentation and has been used as a target in previous studies of this species. We attempted to use the same sgRNAs as that in previous studies (sgRNA460 and sgRNA519* in Basu et al., 2015 and Chaverra-Rodriguez et al., 2018); however we found that there was a SNP in the target site of sgRNA519* in our strain (data not shown). Therefore, we used two sgRNAs: one was newly designed (sgRNA1) and the other was identical to sgRNA460 (Basu et al. 2015; Chaverra-Rodriguez et al. 2018) (sgRNA2) (Fig. 1A).

**Fig. 1.**
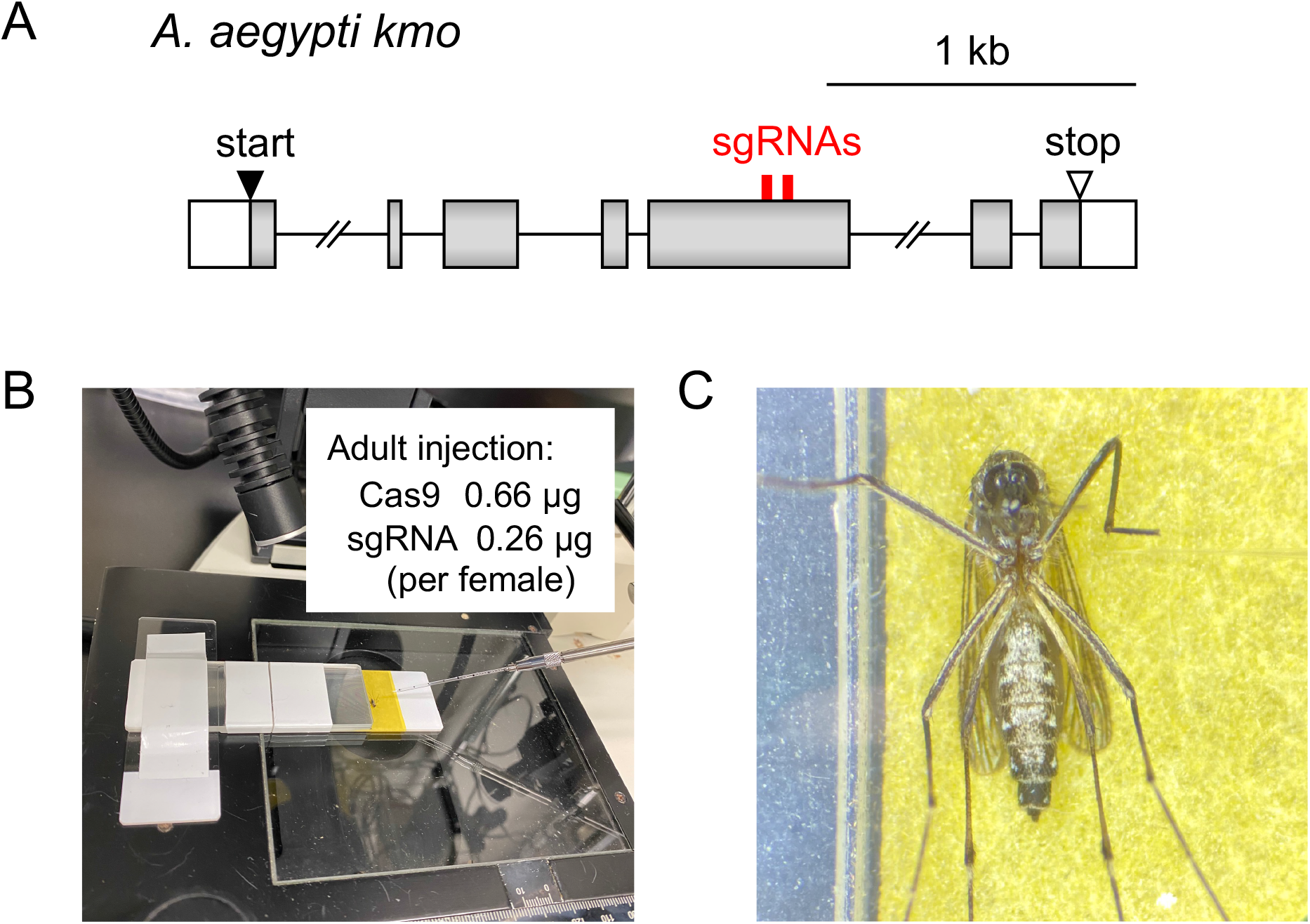
DIPA-CRISPR in *Aedes aegypti*. (A) DIPA-CRISPR target sites of the *A. aegypti kmo* gene (NC_035108.1). The red box indicates the sgRNA sites. (B) Adult injection in *A. aegypti*. (C) Enlarged images of injected *A. aegypti*. Females were injected intrathoracically.

In *A. aegypti*, synchronous egg development is regulated by blood-feeding (Raikhel and Dhadialla 1992). Because the highest gene editing efficiency was achieved by injecting females at the vitellogenic stage in *B. germanica* and *T. castaneum* (Shirai et al., 2022), we evaluated three time points around the onset of vitellogenesis in *A. aegypti*, specifically, 6, 24, and 48 h after blood-feeding (ABF).

### DIPA-CRISPR experiments in *A. aegypti*

We prepared a mixture of commercial Cas9 and the two sgRNAs targeting *kmo* and injected it into females at selected hours ABF (Fig. 1B, C). When we screened the eye colors of the progeny (generation zero, G_0_), we were able to recover a white-eyed G_0_ individual from the progeny of females injected at 24 h ABF (Fig. 2A). To verify that the desired gene editing events had occurred in this white-eyed animal, we performed genomic PCR and observed two DNA bands, which indicated the presence of a large deletion allele (Fig. 2B). Sanger sequencing of the subcloned PCR products revealed that there were multiple types of edited alleles in this white-eyed animal, including the large deletion allele (Fig. 2C). Of note, all 15 sequenced clones carried insertions and/or deletions; thus, it is likely that almost all cells of this G_0_ white mutant carried edited alleles. Because *kmo* is an autosomal gene, the recovery of the white-eyed animal indicates that Cas9 RNPs induced biallelic (i.e., both maternal and paternal) mutations in this animal.

**Fig. 2.**
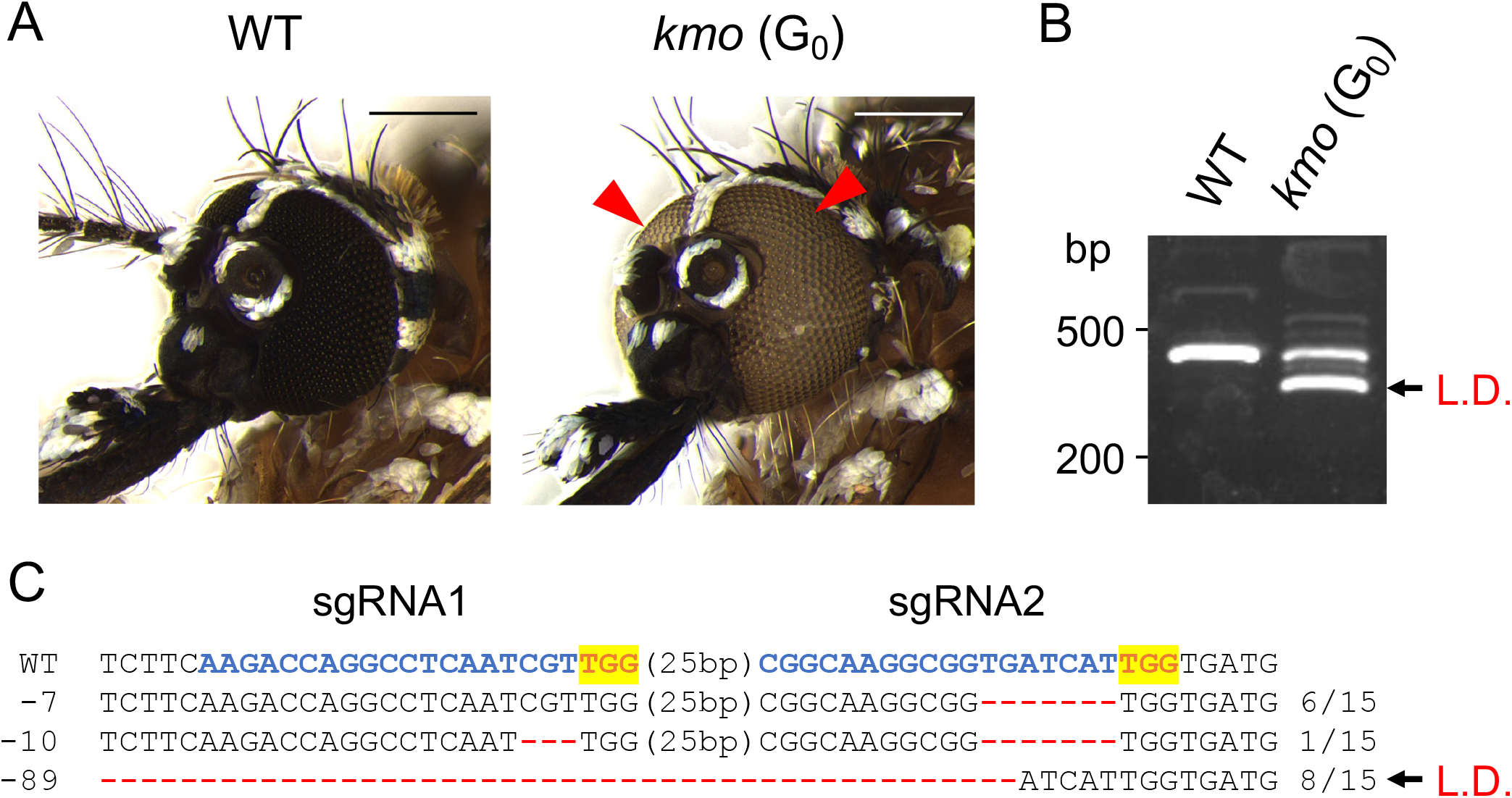
A white-eyed G_0_ individual. (A) A recovered white-eyed G_0_ adult. Arrowheads indicate white eyes. Scale bars, 250 μm. (B) Detection of a large genomic deletion in the white-eyed G_0_ animal by genomic PCR. (C) The DNA sequences of the edited alleles in the white-eyed G_0_ animal. Blue letters represent sgRNAs and highlighted orange letters indicate PAM sequences. L.D. indicates the induced large deletion.

### Screening of edited animals by genotyping

To further screen for mosquitoes with edited alleles without detectable external phenotypes, 288 G_0_ individuals from each experimental condition were randomly selected and subjected to HMA. We obtained one edited animal from the batch injected at 6 h ABF (GEF = 0.3%) and 10 animals from the batch injected at 24 h ABF, respectively (GEF = 3.5%) (Fig. 3A and Table 1). In contrast, we were unable to recover any gene-edited animals from the batch injected at 48 h ABF (Table 1).

**Fig 3.**
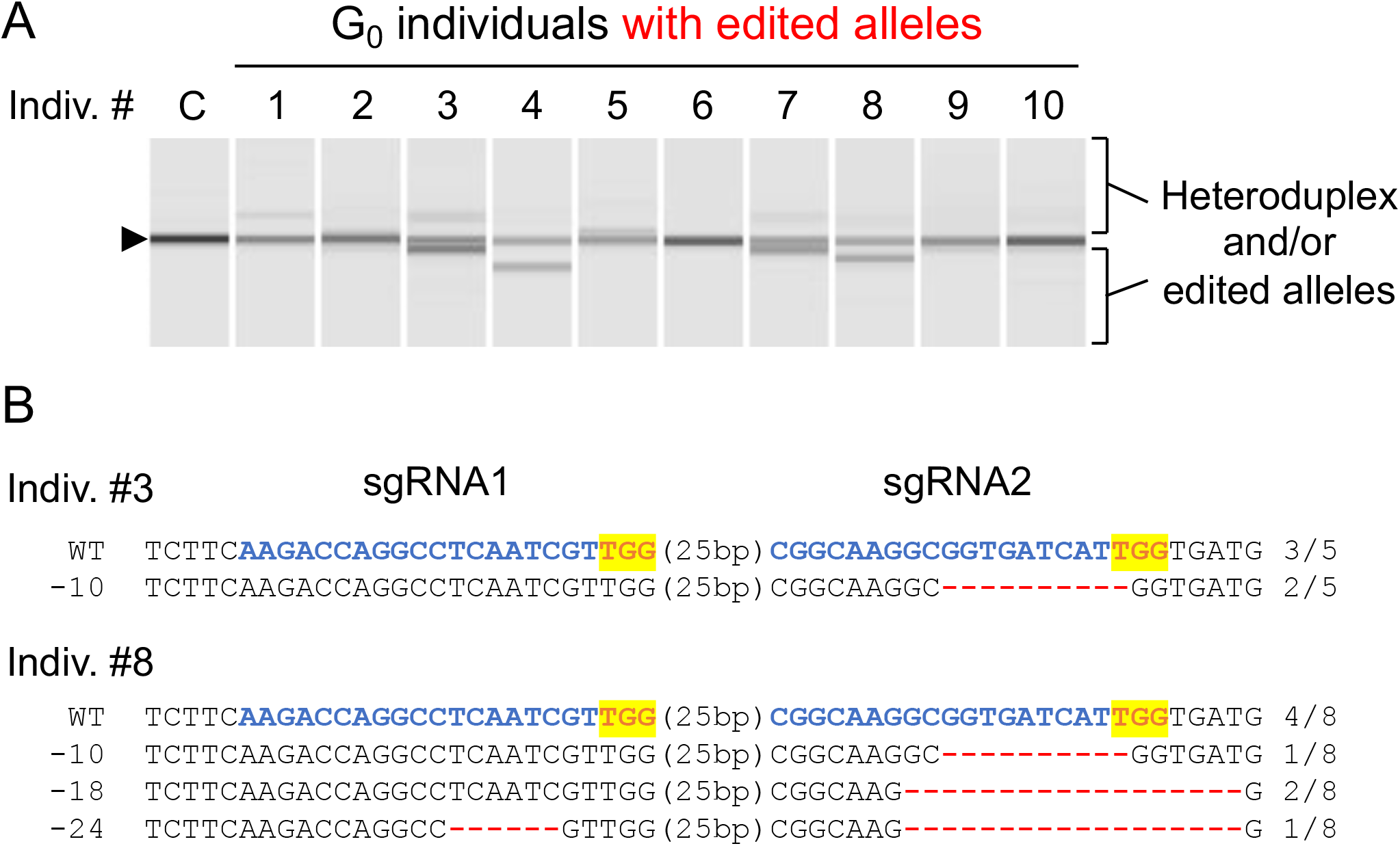
The results of genotyping in G_0_ individuals. (A) The results of the heteroduplex mobility assay (HMA) in G_0_ individuals carrying edited alleles. An arrowhead indicates the band from the wild-type allele. (B) The DNA sequences of the mutant alleles from individuals #3 and #8. Blue letters indicate sgRNAs and highlighted orange letters indicate PAM sequences.

**Table 1.** The efficiency of DIPA-CRISPR in *Aedes aegypti*.

| Protein | Time of<br>injection ABF<sup>a</sup> | Injected<br>females | Survival<br>rates<sup>b</sup> (%) | Screened G<sub>0</sub><br>by genotyping | G<sub>0</sub> with<br>edited alleles | GEF<sup>c</sup> (%) |
| --- | --- | --- | --- | --- | --- | --- |
| Cas9 | 6 h ABF | 40 | 82.5% | 288 | 1 | 0.3% |
| (IDT) | 24 h ABF | 40 | 75% | 288 | 10 | 3.5% |
|  | 48 h ABF | 41 | 100% | 288 | 0 | 0% |

a, ABF, after blood-feeding
b, Survival rates were calculated between 1–3 days post injection.
c, GEF, gene editing efficiency

To determine the nucleotide sequences of the edited alleles, two individuals (#3 and #8 in Fig. 3A) were subjected to Sanger sequencing. For individual #3, two of the five subclones contained a 10-bp deletion at the site of sgRNA2 (Fig. 3B). In the eight subclones from individual #8, one contained a 10-bp deletion, two had an 18-bp deletion, and one harbored a 24-bp deletion (Fig. 3B). These results indicate that both sgRNAs exhibited gene editing activity and that multiple gene editing events occurred in the oocytes before and/or after fertilization. Importantly, we show that the injection at 24 h ABF exhibited the highest gene editing efficiency (GEF = 3.5%) (Table 1), which corresponds to the time of vitellogenesis (Koller et al. 1989; Koller and Raikhel 1991).

## Discussion

In the present study, we successfully adapted DIPA-CRISPR to the yellow fever mosquito *A. aegypti*, which contains the derived polytrophic meroistic ovaries. We also found that the 24 h ABF injection exhibited the highest gene editing efficiency (Table 1). In *A. aegypti*, the uptake of yolk protein precursors peaks between 24 and 30 h ABF, followed by a rapid decline and a cessation of uptake by 36 h (Koller et al. 1989; Koller and Raikhel 1991). Therefore, the peak of GEF values corresponds well with the time of the vitellogenesis in *A. aegypti*. Similar results were obtained in *B. germanica* and *T. castaneum*, in which the highest GEF values were achieved in females undergoing vitellogenesis. Therefore, our results clearly demonstrate that a key parameter for DIPA-CRISPR is the stage at which the females injected.

There is another method of gene editing by adult injection, known as Receptor-Mediated Ovary Transduction of Cargo (ReMOT) (Chaverra-Rodriguez et al. 2018). Thus far, ReMOT has been applied to several species, including mosquitoes (Chaverra-Rodriguez et al. 2018; Macias et al. 2020; Li et al. 2021) wasps (Chaverra-Rodriguez et al. 2020), beetles (Shirai and Daimon 2020), whiteflies (Heu et al. 2020), and a non-insect tick species (Sharma et al. 2022). However, this method has some limitations, such as the need to produce recombinant Cas9 protein fused to peptide ligands. This facilitates ovary transduction and the use of endosomal escape reagents (EERs) that are thought to facilitate the release of Cas9 RNPs from the endosome into the cytosol. In the previous ReMOT study, the same target gene *kmo* was disrupted in the same species (Chaverra-Rodriguez et al. 2018) and the highest GEF was 2.4%. Although we cannot directly compare the efficiencies of two different studies, we conclude that the efficiency of our DIPA-CRISPR approach (GEF = 3.5%) was comparable to that of ReMOT. Because of the simplicity of the procedure, which eliminates the need for engineering, preparation of recombinant Cas9 proteins, and the use and optimization of EERs, DIPA-CRISPR is a more feasible and accessible approach for gene editing in mosquitoes.

We previously demonstrated that DIPA-CRISPR enables efficient gene editing in insects with panositic (*B. germanica*) and telotrophic (*T. castaneum*) ovaries. Therefore, our successful gene editing in mosquitoes using DIPA-CRISPR strongly suggests that this approach can be applied to all three types of ovaries. In addition, gene editing was achieved in the spider mite (Dermauw et al. 2020) by adult injection of non-tagged Cas9 and sgRNA. We anticipate that DIPA-CRISPR will be extended to a wide variety of non-model insects and other arthropods as a research tool for answering fundamental biological questions as well as for the control of agricultural and medical pests.

## Acknowledgments

We thank Takahiro Ohde, Toshiya Ando, and the laboratory members for helpful suggestions and discussion, and Misaki Masuda for assistance with mosquito rearing. This research was supported by JSPS KAKENHI (nos. 20K21311and 22K19179 to T.D.) and Moonshot Research and Development Program for Agriculture, Forestry, and Fisheries (Funding Agency: Bio-oriented Technology Research Advancement Institution) (no. JPJ009237 to T.D. and M.O.). Y.S. is a JSPS Research Fellow with a research grant (DC1, 21J20658).

